# Sensitive and robust assessment of ChIP-seq read distribution using a strand-shift profile

**DOI:** 10.1101/165050

**Authors:** Ryuichiro Nakato, Katsuhiko Shirahige

**Affiliations:** Institute of Molecular and Cellular Biosciences, The University of Tokyo, 1-1-1 Yayoi, Bunkyo-ku, Tokyo, 113-0032, Japan

## Abstract

**Abstract:** Chromatin immunoprecipitation followed by sequencing (ChIP-seq) can detect read-enriched DNA loci for point-source (e.g., transcription factor binding) and broad-source factors (e.g., various histone modifications). Although numerous quality metrics for ChIP-seq data have been developed, the ‘peaks’ thus obtained are still difficult to assess with respect to signal-to-noise ratio (S/N) and the percentage of false positives.

We developed a quality-assessment tool for ChIP-seq data, SSP (strand-shift profile), that quantifies S/N and peak reliability without peak calling. We validated SSP in-depth using ≥ 1,000 publicly available ChIP-seq datasets along with virtual data to demonstrate that SSP is quantifiable and sensitive to different S/Ns for both point- and broad-source factors. Moreover, SSP is consistent among cell types and with respect to variance of sequencing depth, and identifies low-quality samples that cannot be identified by quality metrics currently available. Finally, we show that “hidden-duplicate reads” cause aberrantly high S/Ns, and SSP provides an additional metric to avoid them, which can also contribute to estimation of peak mode (point- or broad-source) of samples.

**Availability:** https://github.com/rnakato/SSP"

## 1 Introduction

Chromatin immunoprecipitation followed by sequencing (ChIP-seq) can identify DNA loci bound by transcriptional factors (i.e., point-source) as well as broadly distributed histone modifications (i.e., broad-source) [1, 2]. In a ChIP experiment, immunoprecipitated DNA fragments are sequenced as reads that are mapped to a reference genome, and statistically significant read enrichments (as compared with a corresponding Input sample) are detected as peaks. Because technical biases in read distribution may hinder obtaining biologically relevant results [3], quality assessment of ChIP-seq data is critical to ensure that they are of high quality and suitable for subsequent analyses. Numerous computational measures have been developed such as library complexity and GC content [4, 5]. However, quality metrics currently available cannot identify all types of low-quality samples.

The signal-to-noise ratio (S/N) in ChIP-seq assesses the number of peaks over the whole genome, and the value should be high for ChIP samples and low for Input samples. A low S/N for ChIP samples indicates the failure of the immunoprecipitation step, whereas a high S/N for Input samples reflects higher than expected levels of read clustering—both of which should be avoided. A straightforward way to evaluate the S/N is to calculate the fraction of reads falling within peak regions (called FRiP), but it depends on peak-calling parameters and sequencing depth (hereafter we use “depth” for simplicity). In contrast, cross-correlation analysis [4] evaluates the S/N without the need for peak calling. A crosscorrelation analysis estimates the Pearson correlation coefficient between the read densities mapped on the forward and reverse DNA strands upon shifting from one strand to the other (see Fig. S1 for an example). Such a “strand-shift profile” typically peaks at the shift corresponding to the DNA fragment length, which increases with increasing S/N for the sample. This tendency has also been used to estimate fragment length from single-end reads. There is also a spike at the read-length shift that arises from repetitive sequences [6]. Based on this observation, crosscorrelation analysis calculates two metrics, namely the normalized strand coefficient (NSC) and the relative strand correlation (RSC, see subsection 2.1 for details). These metrics have been used tin large con sortia such as ENCODE [7] and ROADMAP [8]. A strand-shift profile strategy based on the Hamming distance was also proposed for rapid computation [9]. Whereas these tools are useful for point-source factors, broad-source factors (e.g., H3K9me3) often have marginal or truly low scores compared with Input samples owing to the relatively low peak intensity (height), even when the samples are of high quality [4]. Moreover, these S/N indicators do not evaluate the reliability of the peaks obtained, that is, the percentage of false positives derived from read distribution bias (e.g., GC bias) [5]. Visual inspection of a limited number of sites is effective but insufficient to explain the properties of read distribution over the whole genome. Consequently, genome-wide assessment of peak quality still presents challenges that current protocols cannot circumvent.

To address this shortcoming, we developed SSP, which is based on a strand-shift profile using the Jaccard index to assess S/N and read distribution bias in ChIP-seq datasets. We evaluated the performance of SSP using an extensive dataset of ChIP-seq samples for various cell types obtained from the ENCODE and ROADMAP projects, along with virtual data, to demonstrate that SSP is quantifiable and sensitive to different S/Ns for both point- and broad-source factors and is consistent among cell types and different depths. We also found that RSC is inappropriate for calculating the S/N. Finally, we show that “hidden-duplicate reads” confound the strand-shift profile because they cause unexpected enrichment, resulting in aberrantly high S/Ns (see section 3.6.1). Therefore we developed an additional metric to overcome this problem, which can also be used to estimate peak mode (point or broad source) without peak calling.

## 2 Methods

### 2.1 Strand-shift profile using the Jaccard index

Figure 1 presents an overview of SSP. Using mapped reads as input, SSP generates the strand-specific vectors for forward and reverse strands of each chromosome *c* 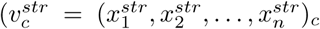 *str ∈ {fwd, rev },* step 1) Because reads mapped to the same genomic position are removed as duplicate reads [4], each element of a strand-specific vector is binary—either zero (unmapped) or one (mapped).

**Figure 1:**
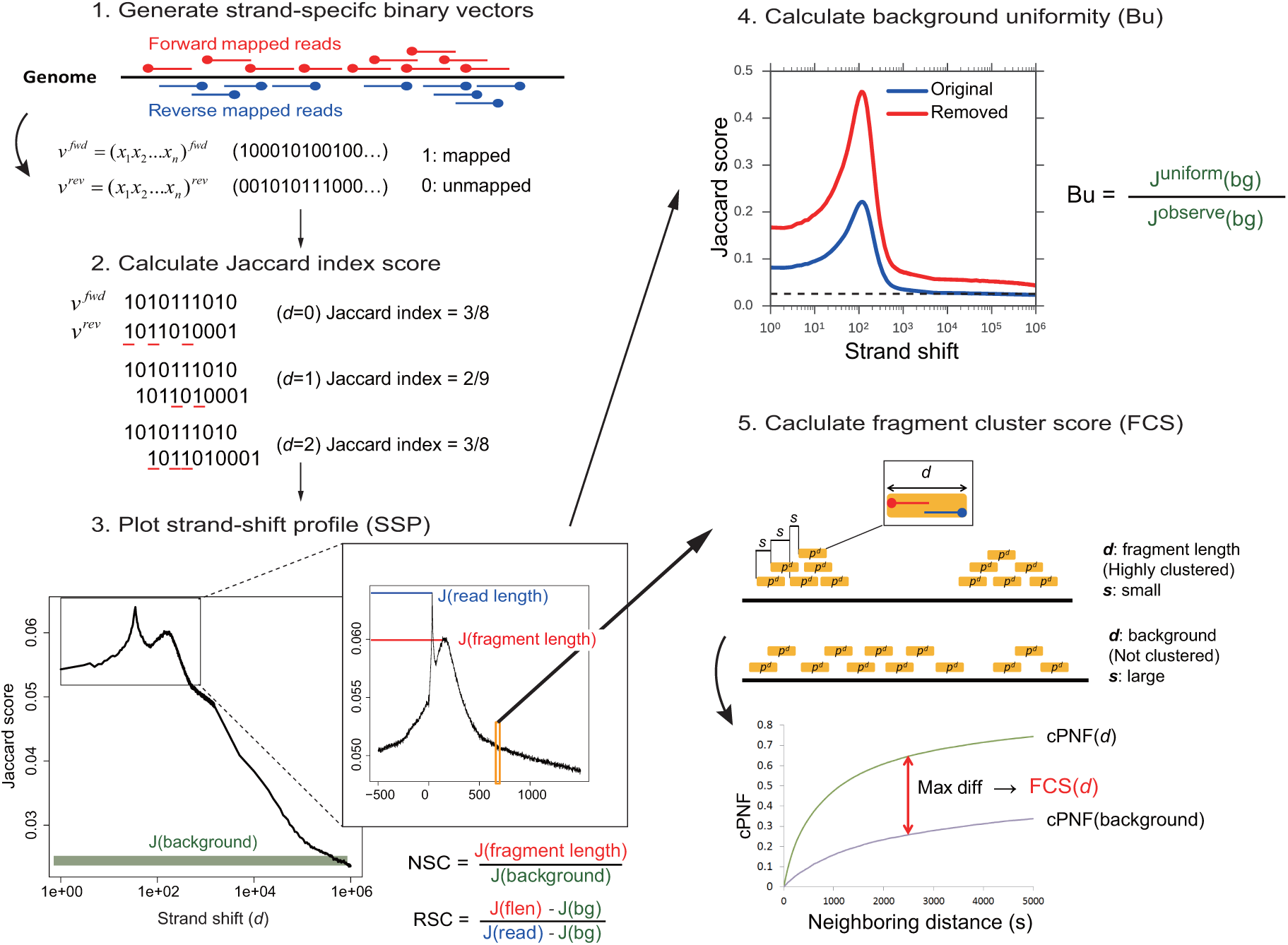
Workflow for SSP. Step 1: convert mapped reads to strand-specific binary vectors (*n*: chromosome length), in which ‘1’ indicates that the 5’ end of a read is mapped at the genomic position. Duplicate reads are discarded. Step 2: calculate the similarity between forward and reverse strand-specific binary vectors for each strand shift *d* based on the Jaccard index. An example calculation is shown (*n* = 10, *d* = 0, 1, 2). Step 3: plot a strand-shift profile based on the Jaccard index and calculate NSC and RSC. Fragment length is estimated as the distance *d* at which the Jaccard score is maximal except for read-length shift. Step 4: calculate background uniformity based on the background level. A strand-shift profile of GM12878 PU.1 data from ENCODE is shown. Blue, original data. Red, data after removing mapped reads in every other 10-Mbp window. The horizontal dashed line indicates the expected background level. Step 5: calculate the fragment cluster score to evaluate the cluster level of all forward-reverse read pairs with each distance *d* (orange rectangles), where *s* is the distance to the nearest downstream read pair. These read pairs are the same as the red bars in step 2. cPNF is the cumulative proportion of neighboring downstream fragments.

SSP calculates the Jaccard index between 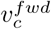 and 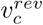 for each strand shift *d* (-500 bp *< d <* 1 Mbp) as follows (step 2):

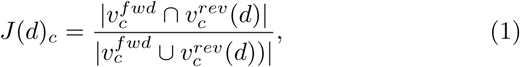

where 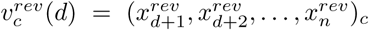 Whereas the Pearson correlation and Hamming distance confer equal weight to pairs of mapped bases (1,1) and unmapped bases (0,0), SSP adopts the Jaccard index that focuses on the mapped bases (1,1) because unmapped bases can often coincide as a consequence of the lack of depth and low-mappable regions.

To standardize the value for different genome lengths, the Jaccard score is normalized by the number of mapped reads (*N*_*c*_) and the number of uniquely mappable positions on chromosome *c* (*L*_*c*_):

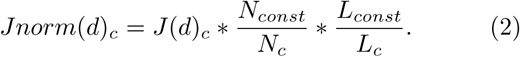

*N*_*const*_ and *L*_*const*_ are arbitrary constants (default, 10 million and 100 million). SSP assembles Jaccard index profiles for all autosomes (step 3):

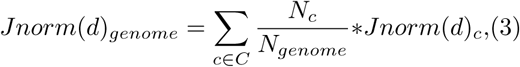

where *C* is the set of all autosomes and *N*_*genome*_ *= ∑*_*c∈C*_ *N*^*c*^ Sex chromosomes are excluded to ignore sex-specific differences. We use this *Jnorm*(*d*)_*genome*_as Jaccard score *J* (*d*) for each sample in SSP. The magnitude of *Jnorm*(*d*) reflects the co-occurrence of reads mapped on the forward and reverse strands with distance *d*.

The fragment length *d*_*flen*_is estimated as *argmaxJ* (*d*). NSC and RSC are then calculated in the same manner as a cross-correlation analysis, that is, *NSC* = *J* (*d*_*flen*_)*/J* (*d*_*bg*_) and *RSC* = (*J* (*d*_*flen*_) *-J* (*d*_*bg*_))*/*(*J* (*d*_*readlen*_) − *J* (*d*_*bg*_)), where *J* (*d*_*bg*_) is the Jaccard score for the background. Whereas existing methods use *d* = 1,000–1,500 bp as background, SSP takes the average over a range of 500 kbp to 1 Mbp at steps of 5 kbp (default) because we observed that the Jaccard score still decreases up to 1 Mbp for the human genome (Fig. 1, step 3).

### 2.2 Background uniformity

A fundamental question is “Why does background level *J* (*d*_*bg*_) vary among samples?” By definition, *J* (*d*_*bg*_) reflects the co-occurrence probability of forward and reverse reads. Ideally, the background reads should be uniformly distributed; in reality, however, the read distribution is often more congregated, or biased, owing to various potential technical or biological issues [3], resulting in a higher *J* (*d*_*bg*_). Although the library complexity evaluates the percentage of duplicate reads, it does not directly reflect any potential bias in the read distribution. In fact, we observed that *J* (*d*_*bg*_) increased ∼2-fold when the mapped reads were removed in every other 10-Mbp window, whereas library complexity and NSC score remained essentially unchanged (Fig. 1, step 4).

Based on this observation, we defined “background uniformity” (Bu, step 4):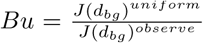, where *J(d*_*bg*_)^*uniform*^ is the Jaccard score of background for a sample that has completely uniform read distribution (see Supplementary Methods for calculating *J(d*_*bg*_)^*uniform*^ A large value for Bu indicates that the sample has a relatively uniform read distribution in background regions. Bu should range from 0 to 1, but practically the maximum score for Bu slightly exceeds 1.0 because the estimated number of *L*_*c*_is slightly larger than it actually is.

### 2.3 Fragment cluster score

Finally, SSP calculates a “fragment cluster score” (FCS) that estimates the cluster level of forward-reverse read pairs with each strand shift *d* (step 5).

Pairs of mapped bases (1,1) with strand shift *d* can be expressed as read pairs mapped on forward and reverse strands with distance *d* (*p*^*d*^, orange rectangles in Figure 1). All read-pairs are denoted 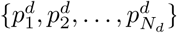, where 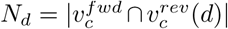, which is sorted by genomic position. Let *f* (*d, s*) represent the number of 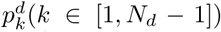 that have neighboring read pairs 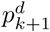 within distance *s*. Then the cumulative proportion of neighboring fragments (cPNF) is:

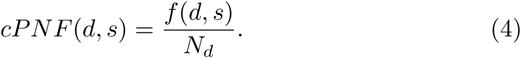

cPNF is calculated up to *s*_*maxs*_(default, 5 kbp). FCS(d) is defined as the maximum difference of cPNF(*d*) from cPNF(*d*_*bg*_) against *s* (Fig. 1):

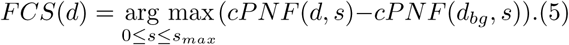

Because FCS(d) depends on the depth, SSP calculates it with a fixed number of reads (default, 10 million reads). This maximum difference strategy provides more robust values against different values for *s*_*max*_and *d*_*bg*_than the relative entropy such as the Kullback-Leibler divergence. The resulting FCS profile reflects the cluster level of *p*^*d*^ in the sample, whereas the Jaccard score *J* (*d*) reflects the number of *p*^*d*^ (i.e., *N*_d_). Figure S2 shows the typical pattern of cPNF and FCS profile. SSP uses the value at fragment length FCS(*d*_*flen*_) as the FCS score for each sample.

### 2.4 Comparison with current methods

To assess the performance of SSP for estimating S/N, we implemented three existing tools: 1) phantompeakqualtools (PPQT), which internally implements spp version 1.14 [10] for cross-correlation analysis and then outputs NSC and RSC; 2) Q version 1.2.0 [9], which adopts a strand-shift profile based on the Hamming distance and calculates RSC; and 3) deepTools version 2.5.0 [11], which computes the synthetic Jensen-Shannon distance (JSD) that evaluates differences in the cumulative fraction of mapped reads between ChIP samples by assuming a Poisson distribution as a background model for windows of fixed length. We applied deepTools with the “ignoreDuplicates” option according to the instructions given in the manual. We used default parameters for each of the other tools.

### 2.5 Other quality scores

FRiP, library complexity for 10 million reads, peak height and GC content distribution of nonredundant reads were calculated with DROMPA version 3.2.6 [12]. “GC peak” refers to the summit position of GC content profile.

## 3 Results

### 3.1 Estimating fragment length

We first evaluated the performance of fragmentlength estimation with SSP, PPQT, and Q using 45 paired-end ChIP-seq datasets for human (Fig. 2A and Table S1, see Supplementary Methods for detailed protocol). We found that SSP provided comparable and more accurate fragment-length data than PPQT and Q. PPQT and Q were nearly as accurate as SSP but could not provide a fragment-length estimate for several of the samples (e.g., samples 37 (GM12878 H3K4me1) and 45 (K562 WCE)). On the other hand, none of the programs could estimate a fragment-length for certain samples (e.g., sample 16 (K562 H3K27me3)) for which there was no clear peak in the strand-shift profile (Fig. S3). It has been reported that a high score for read-length shift can be mitigated by removing reads mapped on “blacklist regions” in the genome [6]. However, removing reads mapped on them had little effect (Fig. S4). In fact, because a failure of fragment-length estimation is mainly attributable to a lack of enrichment at the fragment-length shift, mitigating the enrichment at read-length alone is insufficient. In this case, fragment length should be supplied by the users.

**Figure 2:**
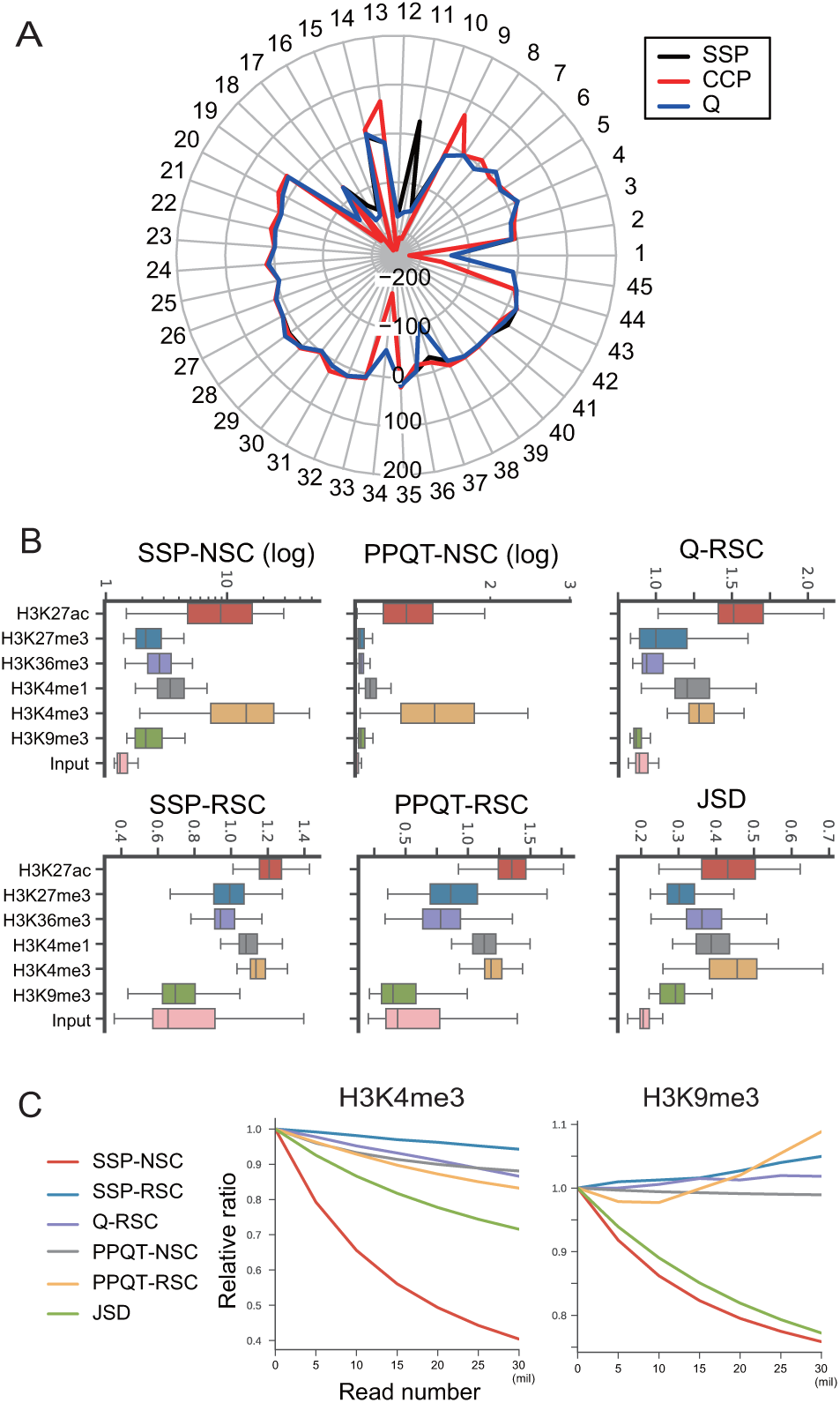
(A) Radar plot of the comparison between the fragment-length estimated by each tool and that from paired-end data for 45 paired-end ChIP-seq data for human (build hg19). The *y* axis indicates the difference between the fragment size estimated from the single-end (forward) reads by these tools and that derived from the paired-end reads. (B) Distribution of scores by SSP (NSC and RSC), PPQT (NSC and RSC), Q (RSC), and deepTools (JSD). Note that the *y* axis is a log-scale for SSP-NSC and PPQT-NSC but a linear scale for the others. (C) Relative ratio at different S/N values (adding Input reads from 5 million to 30 million) against original data.

In subsequent analyses, we did not remove blacklist regions because doing so could affect RSC, and in fact detailed blacklist regions are available only for human genome build hg19.

### 3.2 SSP and JSD archive sufficient sensitivity both for point- and broad-source factors

To comprehensively evaluate the performance of SSP relative to other tools, we first used a compendium of 860 ChIP-seq samples of six core histone modifications for 127 cell types, consisting of point-source (H3K27ac, H3K4me1, H3K4me3) and broad-source factors (H3K27me3, H3K36me3, H3K9me3) along with Input samples, obtained from the ROADMAP web portal (http://egg2.wustl.edu/roadmap/webportal). We acquired consolidated map data, in which reads had been truncated to 36 bp, mapped onto genome build hg19, filtered based on mappable bases of 36-bp reads, and then uniformly down-sampled to a maximum depth of 30 million reads. This avoided the effect derived from different read lengths, parameters for mapping, and mappability. Moreover, these datasets contain certain low-quality data [13], which is preferable for dataset of QC performance evaluation.

Figure 2B shows the comparison (see Table S2 for detailed information of each sample). The results revealed that SSP-NSC and JSD could achieve sufficient sensitivity both for point- and broad-source marks. The smaller difference between point- and broad-source marks for JSD compared with SSP-NSC is perhaps a consequence of score saturation, i.e., given that the maximum value of JSD is 1.0. PPQTNSC showed little difference (only ∼1.1 fold) among three broad marks compared with Input samples, indicative of insensitivity for broad marks. As previously reported, RSCs calculated with all three tools were comparable or lower for H3K9me3 than Input samples, possibly because H3K9me3 is more highly enriched at the read-length shift (*J* (*d*_*readlen*_)) compared with other histone modifications derived from repetitive regions, such as centromeres [14]. RSC amalgamates the magnitude of true peak enrichment and repeat effects, and thus when *J* (*d*_*readlen*_) is high, RSC may be small even when the S/N is sufficiently high.

To further validate the sensitivity of the S/N indicators, we generated virtual data for histone modifications with various S/Ns by adding a fixed number of Input reads in a stepwise manner (see Supplementary Methods for details). The S/N then decreased with increasing numbers of Input reads. Figure 2C and Figure S5 show the comparison for H3K4me3 (point source) and H3K9me3 (broad source) as well as the other four histone marks, respectively (E072, brain inferior temporal lobe cells). In most cases, the values of the indicators decreased with increasing numbers of Input reads. RSC was relatively greater for H3K9me3 because, for this mark, the scores were often lower than those of the Input (Fig. 2B). SSP-NSC had superior or comparable sensitivity to changes in S/N, whereas PPQT-NSC lacked sensitivity for evaluating broad marks.

### 3.3 SSP is robust for various cell types and different depth

The S/N estimation can be affected by multiple factors, such as depth, read length and copy number variations in cancer cell lines [15]. To validate the robustness of the S/N indicators against these factors, we next utilized 399 ChIP-seq samples of transcriptional factors for 20 cell types obtained from the ENCODE project [16], which contain various read lengths (25, 36, and 50) and depths, and investigated whether each S/N indicator could distinguish between ChIP samples and Input samples from them. We acquired fastq files from the Sequence Read Archive under accession number SRP008797, mapped them on genome build hg38 using bowtie version 1.1.2 [17] and used uniquely mapped reads (“-n2 -m1” option). See Table S3 for detailed information on each sample.

Figure 3A and Figure S6 depict the distribution of SSP-NSC and the other S/N indicators, respectively, for 20 cell types. Whereas the number of samples varied among the cell types, we found that SSP-NSC yielded distinct differences between ChIP and Input samples for all cell types. To compare all indicators in this respect, we displayed the median scores for each cell type (Fig. 3B). For SSP-NSC and PPQT-NSC, median values for ChIP and Input samples were consistently different among all cell types, indicating that a universal threshold value applicable to any samples could be defined for these indicators. For example, SSP-NSC ≥ 3.0 may be a good candidate threshold for transcriptional factor ChIP samples. Meanwhile, RSC and JSD could not sufficiently distinguish ChIP and Input samples. Although ChIP samples had larger values than Input samples for each cell type (Fig. S6), the difference between ChIP and Input samples depended on cell type, and therefore it was difficult to determine a uniform threshold value.

**Figure 3:**
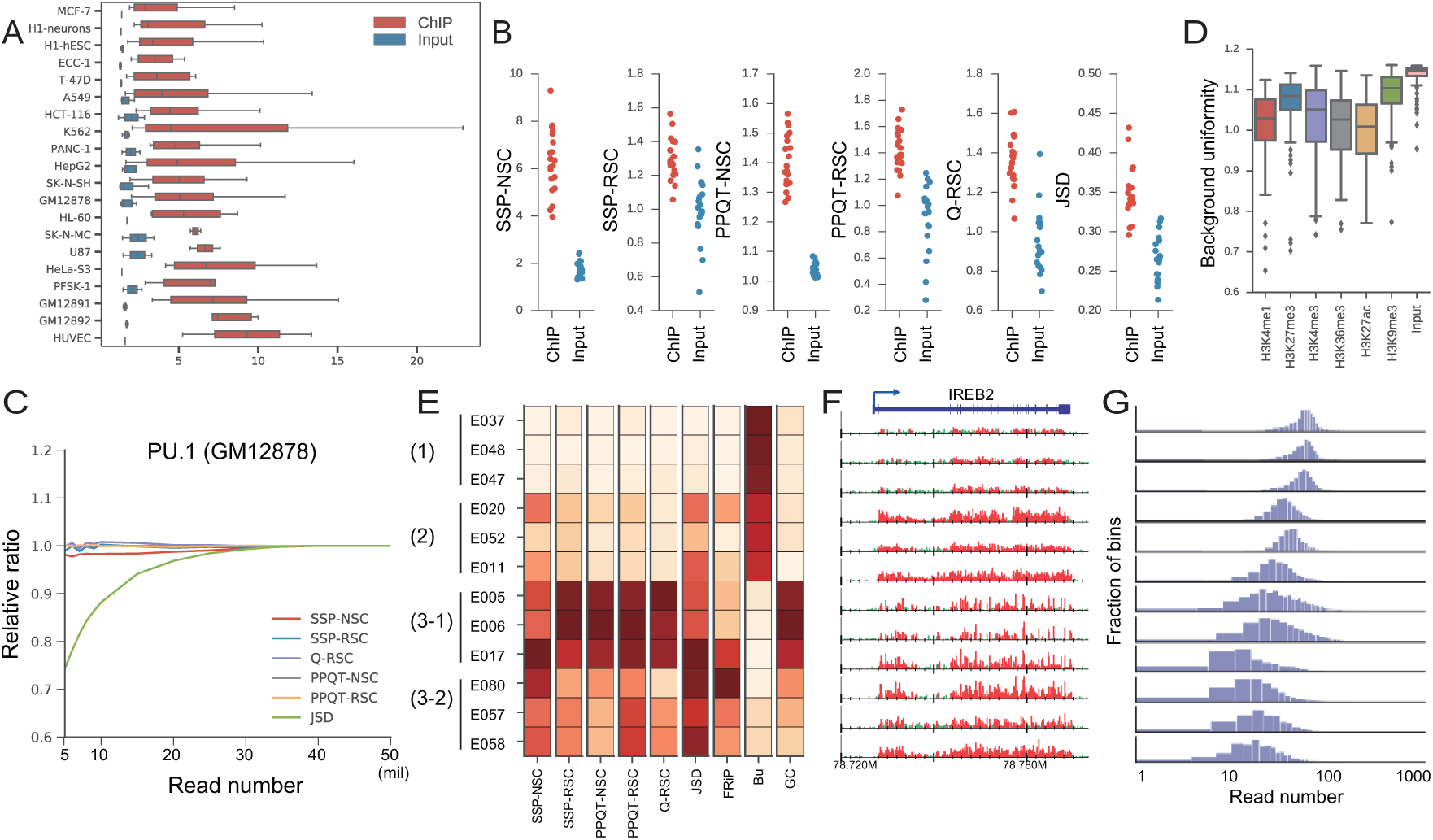
(A) SSP-NSC distribution of ChIP (red) and Input samples (blue) for 20 cell types. (B) Median values for S/N indicators for 20 cell types. (C) Relative S/Ns at each depth (5–50 million) against 50 million reads. Duplicate reads were removed in advance. (D) Distribution of background uniformity for histone modifications. (E–G) Analysis of H3K36me3 data for 12 cell types from ROADMAP. (E) Heatmap of S/N scores alongside Bu and GC peak. Darker colors indicate higher values. (F) Read distribution near the *IREB2* locus (chromosome 15, 78.72–78.80 Mbp). Read number was normalized by the total number of nonredundant reads. The peak regions identified by MACS2 are highlighted in red. (G) Histogram of mapped read number for each 100-kbp bin of the whole genome except chromosome Y.

### 3.4 SSP yields the highest correlation with normalized FRiP, whereas JSD depends on depth

To further evaluate the performance of the S/N indicators, we calculated the Spearman’s correlation between the FRiP and each S/N indicator across the ENCODE and ROADMAP datasets (Table 1). Because the number of obtained peaks for calculating FRiP depends on depth, we calculated two types of FRiP: FRiP based on peaks obtained with total read normalization (FRiPnorm) and without normalization (FRiPraw). FRiPraw is based on peaks identified by MACS2 version 2.1.1 [18], which does not utilize total read normalization (–nomodel option, we also supplied the –broad option for broad marks), and FRiPnorm is based on peaks identified by DROMPA version 3.2.6 (“-n GR” option), which utilizes total read normalization.

**Table 1:** Spearmans correlation between each S/N and FRiP without read normalization (FRiPraw, top values) and with normalization (FRiPnorm, bottom values). *p-value for correlation coefficient *<* 0.01.

|  | SSP-NSC | SSP-RSC | PPQT-NSC | PPQT-RSC | Q-RSC | JSD |
| --- | --- | --- | --- | --- | --- | --- |
| ENCODE | *0.84 | 0.10 | *0.83 | *0.15 | *0.16 | *0.93 |
|  | *0.90 | *0.28 | *0.88 | *0.31 | *0.32 | *0.81 |
| ROADMAP<br>(point source) | *0.77 | *0.20 | *0.70 | *0.33 | *0.29 | *0.97 |
|  | *0.94 | *0.28 | *0.89 | *0.34 | *0.30 | *0.90 |
| ROADMAP<br>(broad source) | *0.72 | *0.40 | *0.43 | *0.37 | *0.37 | *0.90 |
|  | *0.89 | *0.31 | *0.64 | *0.32 | *0.31 | *0.79 |

RSC yielded a low correlation, suggesting that RSC cannot be used for quantification of the S/N. SSP-NSC, PPQT-NSC, and JSD were highly correlated with both types of FRiP. The lesser correlation of PPQT-NSC with broad-source marks compared with point-source marks implies its lower sensitivity for broad-source marks. Whereas SSP-NSC and PPQT-NSC each correlated better with FRiPnorm, JSD correlated more closely with FRiPraw, which clearly shows the dependency of JSD on depth.

To investigate this tendency, we implemented a down-sampling analysis (Fig. 3C and Fig. S7A). We selected three samples that contained an abundant number of reads (*>* 50 million) after removing duplicate reads. For each sample, we subsampled the reads to a fixed number (from 5 million to 50 million) and calculated the ratio of the score at each depth relative to the score for the 50 million reads. No indicators except for JSD fluctuated with depth; JSD decreased at lower depth. The analysis of histone modification data also yielded the same conclusion (Fig. S7B).

Consequently, SSP-NSC is a sensitive and robust S/N estimator for both point-source and broad-source marks, which can be standardized across diverse cell types, read lengths and depths.

### 3.5 Background uniformity identifies low-quality samples

We next computed Bu scores for 860 histone modification samples from ROADMAP (Fig. 3D and Table S2). Although these consolidated data did not contain duplicate reads, we noted that a small amount of data had a low Bu score (*<* 0.8). To investigate the various aspects of Bu, we chose 12 H3K36me3 samples as representatives, and the results are shown in Figure 3E–G. We grouped these samples into four types: (1) low NSC and high Bu, (2) high NSC and high Bu, (3) high NSC and low Bu; this latter type was further classified as 3-1 (GC-rich) and 3-2 (not GC-rich). Figure 3F depicts the read distribution proximal to the housekeeping gene *IREB2* [19]. Groups 2 and 3 had high S/Ns, reflecting read enrichment at the *IREB2* locus. Group 3-1, however, had an unexpectedly sparse read distribution, which was not expected considering that H3K36me3 is broadly distributed within genic regions. Considering the GC-richness, this read distribution may be a consequence of GC bias [20]. Interestingly, this group had a striking peak at fragmentlength shift in the strand-shift profile (Fig. S8). This phenomenon might also facilitate the identification of read-distribution bias.

In contrast, group 3-2 had low Bu values without GC bias, and read distribution was more in line with expectations compared with group 3-1. However, this group also had lower overall genome coverage (Fig. 3G). A possible reason for this is that the DNA fragmentation of tightly packed regions, e.g., heterochromatin, did not work well, resulting in a much lower number of reads on the regions. Such a sample might confound the data normalization for comparative analyses that assume comparable read depth among samples [21].

Consequently, Bu is an effective criterion with which to judge whether a specific consideration or a rejection is required for comparative analyses.

### 3.6 FCS can identify peak intensity and peak mode

#### 3.6.1 Hidden duplicate reads and FCS

While verifying the effectiveness of SSP-NSC for calculating the S/N, we also found that strand-shift profiles of a small number of Input samples had peaks at fragment length despite having a low FRiP (e.g., Input of E024 and E058 cells, Figure 4A). These two samples in particular had extremely high SSP-RSC values (6.66 and 5.35), a phenomenon that is commonly observed with PPQT and Q (Fig. S9). We presumed that this is attributable to hidden duplicate reads—that is, at most two reads (forward and reverse pair) that are derived from the same amplified DNA fragment can remain after duplicate-reads filtering because forward and reverse strands are scanned separately for single-end reads (Fig. 4B) [3]. Such reads may often appear in low-library complexity samples and introduce a spike at the fragment length regardless of whether they are clustered, resulting in aberrant NSC and RSC values. To test this hypothesis, we generated strand-shift profiles for a paired-end sample in which both forward and reverse reads were mapped as ‘single-end’. As expected, the resulting profile showed a remarkable peak at the fragment length shift (Fig. 4C). While NSC increased less drastically (1.53 to 2.54), RSC increased by *>* 3fold (0.61 to 2.29). This result suggested the presence of artifactual S/N enrichment without real peaks in a strand-shift profile, which especially influenced the calculation of RSC.

**Figure 4:**
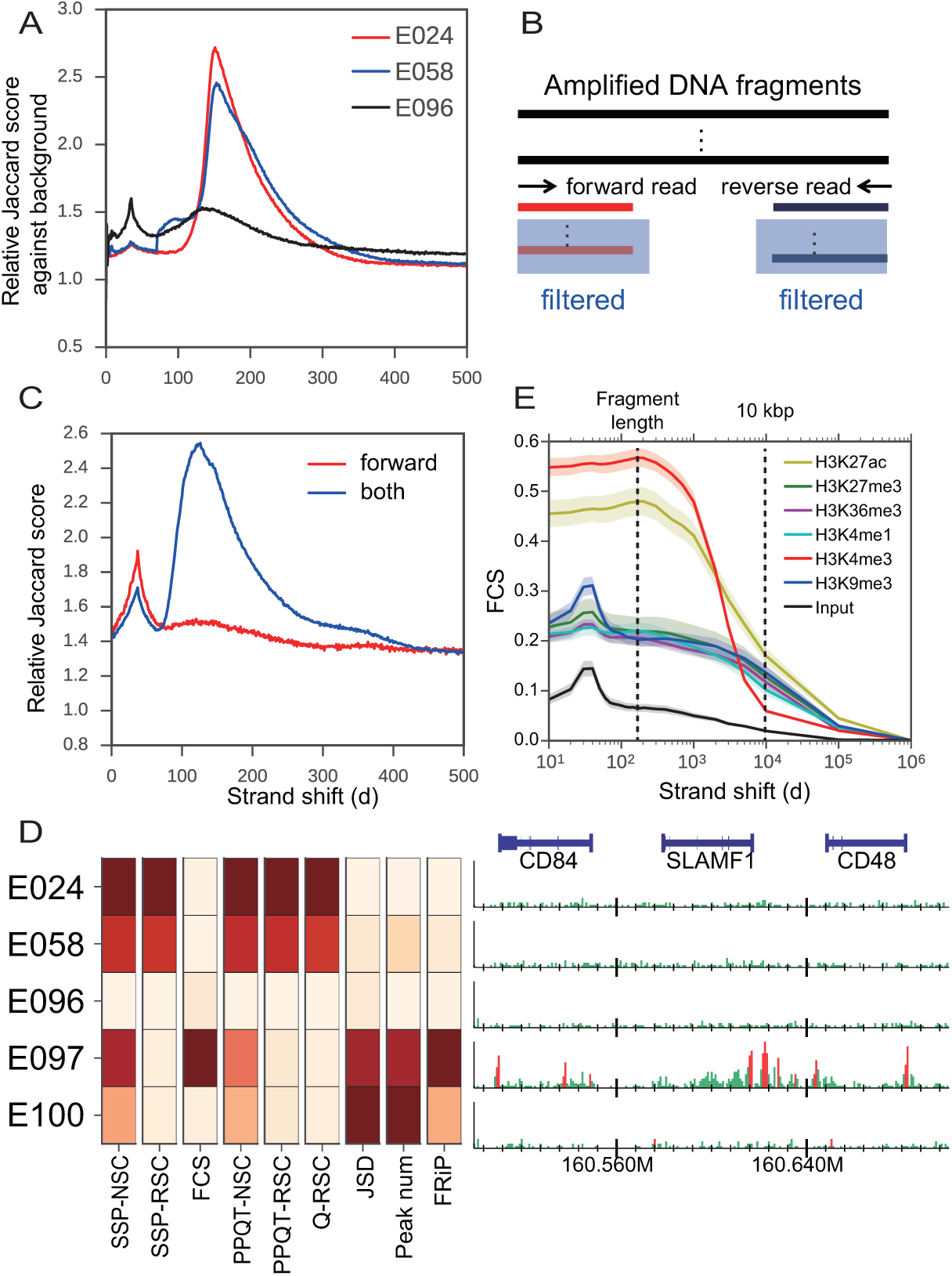
(A) Strand-shift profiles for two Input samples (E024, E058), which have apparent peaks at fragment length but actually have a low FRiP value (*<* 0.01). A typical profile for Input (E096) that has a similar FRiP is also shown. (B) Schematic illustration of hidden duplicate reads. (C) Strand-shift profile for sample K562 H3K9me3 (no. 25 in Figure 2A) using forward read only (red) and both forward and reverse reads as single-end (blue). (D) Heatmap for each S/N value and number of peaks identified by MACS2 for five Input samples alongside their read distributions in the same manner as in Figure 3F (chromosome 1, 160.5–160.7 Mbp). (E) Averaged FCS profiles for histone modification data. Lines and shaded regions indicate the mean value and 95% confidence interval, respectively.

To overcome this problem, we defined FCS, which directly evaluates the cluster level of forward-reverse read pairs with distance *d* (*p*^*d*^, see Methods for details). As the FCS value is high only when *p*^*d*^s are highly clustered as peaks, samples that contain hidden duplicate reads that are not clustered in a genome should have a low FCS score. As expected, FCS could identify read clustering in samples and was little affected by hidden duplicate reads (Fig. S10).

#### 3.6.2 FCS estimates peak intensity

Through our experiments, we found that FCS correlated better with peak intensity than did FRiP, which represents a composite of peak number and intensity (Fig. S11). Figure 4D illustrates the example of five Input samples from ROADMAP. E097 had strong peaks, reflected by the highest FCS score (0.240), which probably has to be rejected from further analysis. In contrast, E100 had more peaks (33,476) than E097, but inspection of the read distribution and relatively lower FRiP suggested that E100 had only small peaks, resulting in a low FCS score (0.044). Therefore, at a sufficiently high peakcalling threshold, most of the small peaks (i.e., as in E100) would be expected to disappear, in contrast to the expectation for E097. E024 and E058 (shown in Figure 4A) had high NSC and RSC values without many peaks, resulting in a low FCS score (0.041 and 0.038, respectively). Although JSD was also minimally affected by hidden duplicate reads because it is not based on a strand-shift profile, it provided E100 with the highest score, suggesting that it correlated better with peak number than did peak intensity and FRiP.

#### 3.6.3 FCS profile has the potential to identify peak mode

Interestingly, a FCS profile that estimates peak intensity with each strand shift *d* reflects peak mode (point or broad source) of histone modifications (Fig. 4E). H3K4me3 had the highest FCS at *d* = fragment length and decreased steeply at *d ≥* 10 kbp. The broad-source marks H3K27me3, H3K36me3, and H3K9me3 each had a moderate score at fragment length, and the value was retained even at *d ≥* 10 kbp, resulting in a higher score than for H3K4me3 at *d* = 10 kbp. H3K27ac had a high score at fragment length and also the highest score at 10 kbp. This is not surprising because H3K27ac had high peaks for point-source marks, some of which clustered in broad genomic regions called super enhancers [22]. This result suggested that FCS has the potential to identify peak mode without the need for peak calling.

## 4. Discussion

To validate whether each sample in a dataset requires special normalization or should be rejected for further analysis, it is crucial to objectively assess the genomewide properties inherent in read distribution. Owing to the difficulty of assessing broad marks, a previous study involving large-scale sample evaluation for S/N was limited to point-source factors and Input samples [23].

Here we introduce SSP, a peak calling-free quality assessment tool for ChIP-seq data. In-depth validation of SSP using ≥ 1,000 publicly available ChIPseq datasets revealed that SSP-NSC achieved sufficient sensitivity and robustness for both point-source and broad-source factors across diverse cell types, read lengths and depths. JSD, as utilized in deepTools, is also sensitive for broad marks, but it has less classification power between ChIP and Input samples among diverse cell types. Moreover, because JSD depends on depth, it requires subsampling for comparison across samples, which is burdensome for large-scale analyses. Although the strand-shift profile strategy (especially RSC) is potentially confounded by hidden-duplicate reads, SSP also provides FCS that avoids their effect. The potential of FCS to evaluate peak mode may facilitate the capture of dynamic changes in genome-wide binding patterns among samples, e.g., in cells during embryonic development [24].

Bu evaluates the reliability of the obtained peaks by quantifying the distribution of mapped reads in background regions. Although GC content correlates with the bias level in ChIP samples, it alone cannot be used for filtering because samples that have many GC-rich peaks (e.g., CpG islands) also have a high GC content. Bu is beneficial in this regard, especially when mapping ratio and library complexity metrics are unavailable (e.g., consolidated datasets).

Although the mean values for these metrics varied among the transcriptional factors and antibodies used (Fig. S12), based on the observations of the datasets in this study and preliminary experiments using several other species (mouse, fly, yeast), we recommend the following thresholds:

- SSP-NSC *≥* 5.0 for strong point-source factors
- SSP-NSC ≥ 1.5 for weak point-source and broad-source factors
- SSP-NSC *≤* 2.0 for Input samples
- FCS(*d*_*flen*_) = 1.5 is the separation between ChIP and Input samples
- Bu *≥* 0.8 for successful ChIP experiments

The exception for Bu was MCF-7 cells, which yielded a relatively lower Bu value possibly owing to extensive copy-number variations (∼0.8, Fig. S12B). The low-Bu samples also were more common when the S/N was extremely high (e.g., RNA polymerase II, Fig. S12C). Thus, it is desirable to use a relaxed threshold value for Bu for these samples. One challenge that remains is to identify falsepositive peaks caused by non-specific binding, such as “hyper-ChIPable regions” [25]. SSP and all existing tools cannot distinguish whether DNA-binding derives from true binding, and thus a comparison with mock ChIP-seq data (e.g., IgG) is needed to avoid such false positives.

## 5 Acknowledgements

We are grateful to Dr. M. Suyama for valuable comments. We would also like to thank our laboratorys members and collaborators.

